# Isoform and protein region abnormalities of dysbindin and copper transporter proteins in postmortem schizophrenia substantia nigra

**DOI:** 10.1101/343178

**Authors:** Kirsten E. Schoonover, Rosalinda C. Roberts

**Affiliations:** Department of Psychology and Behavioral Neuroscience, University of Alabama at Birmingham, Birmingham, AL 35294; Department of Psychiatry and Behavioral Neurobiology, University of Alabama at Birmingham, Birmingham, AL 35294

**Author notes:** Corresponding author information: Kirsten Schoonover, B.A., Sparks Center 866, 1720 2^nd^ Ave South, Birmingham, AL 35294.

**Keywords:** schizophrenia, dysbindin, copper, substantia nigra, postmortem

## Abstract

**Objective:** Dysbindin is downregulated in several schizophrenia brain regions and modulates copper transport required for myelination and monoamine metabolism. We sought to determine dysbindin and copper transporter protein expression in schizophrenia subjects.

**Methods:** We studied the substantia nigra (which exhibits one of the highest copper contents of the human brain) using Western blot analysis. We characterized specific protein domains of copper transporters ATP7A, CTR1, ATP7B, and dysbindin isoforms 1A and 1B/C in postmortem substantia nigra in schizophrenia subjects (n=15) and matched controls (n=11). As a preliminary investigation, we examined medication status in medicated (n=11) versus unmedicated schizophrenia subjects (n=4).

**Results:** The combined schizophrenia group exhibited increased levels of C-terminus, but not N-terminus, ATP7A. Schizophrenia subjects expressed less transmembrane CTR1 and dysbindin 1B/C than controls. When subdivided, the increased C-terminus ATP7A protein was present only in medicated subjects versus controls. Unmedicated subjects exhibited less N-terminus ATP7A protein than controls and medicated subjects, suggesting medication-induced rescue of the ATP7A N-terminus. Transmembrane CTR1 was decreased to a similar extent in both treatment groups versus controls, suggesting no medication effect.

**Conclusions:** These results provide the first evidence of disrupted copper transport into and within schizophrenia nigral cells that may be modulated by specific dysbindin isoforms and antipsychotic treatment.

## Introduction

Schizophrenia manifests in early adulthood with cognitive, positive, and negative symptoms. Several genes have been associated with schizophrenia risk (Schizophrenia Working Group of the Psychiatric Genomics Consortium 2014). The dystrobrevin binding protein 1 (*DTNBP1*) gene encodes the dysbindin protein family, dysbindin-1 (Talbot 2009; Talbot et al. 2009), and is a top candidate gene for schizophrenia (Straub et al. 2002). Allelic variation of *DTNBP1* was associated with schizophrenia shortly after its discovery (Straub et al. 2002), and has been associated with dysbindin protein decreases in cortex, midbrain, and hippocampus in schizophrenia (Talbot 2009; Tang et al. 2009; Weickert et al. 2008; Weickert et al. 2004). *DTNBP1* variations have been associated with impaired white matter integrity in healthy adults (Nickl-Jockschat et al. 2012), loss of grey matter volume in preteenagers (Tognin et al. 2011), abnormalities in neurite outgrowth and morphology (Dickman and Davis 2009), as well as decreased cognitive functions (Burdick et al. 2006), including working, visual, and verbal memory deficits (Varela-Gomez et al. 2015). Additionally, schizophrenia subjects show decreases in specific dysbindin isoforms in a region specific manner, especially in brain regions involved in cognition (e.g., prefrontal cortex and hippocampus) Tang et al. 2009; Weickert et al. 2008; Talbot et al. 2004). Together, these data suggest that dysbindin may be involved in schizophrenia cognitive symptomology, but the mechanism remains unknown.

In mice, dysbindin knockout results in spatial (Cox et al. 2009) and working memory deficits (Papaleo et al. 2012), and increased compulsive and impulsive behaviors (Carr et al. 2013), not unlike the cognitive impairments exhibited by schizophrenia patients with disease-related *DTNBP1* variants (Burdick et al. 2006; Varela-Gomez et al. 2015). Additionally, dysfunctional dysbindin in animal models results in a particularly intriguing consequence of interest: a *decrease* in copper transporters ATP7A and CTR1 (Gokhale et al. 2015). Together ATP7A and CTR1 facilitate copper transport between the blood and the brain (Figure 1A), as well as intracellular transport.

**Figure 1.**
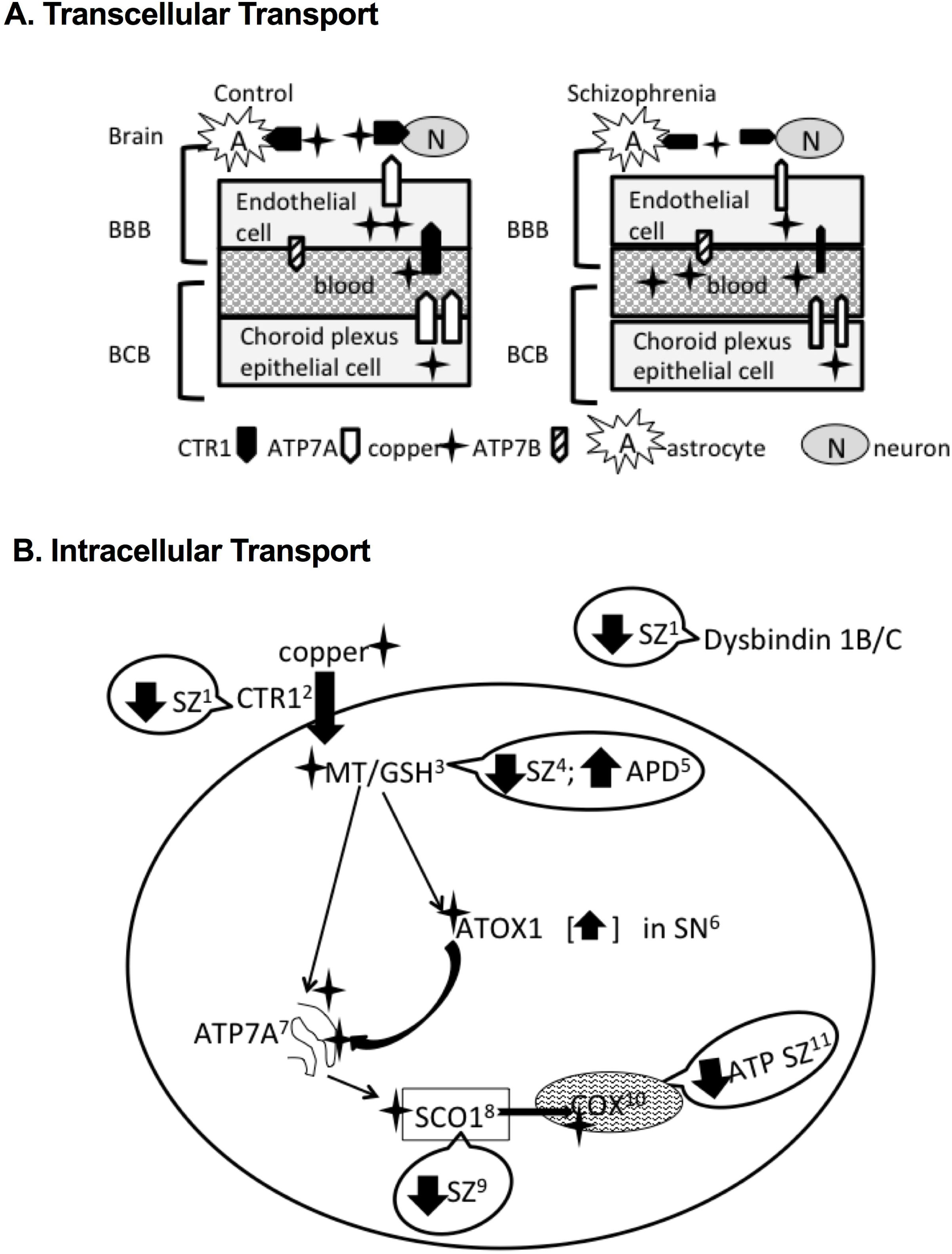
Schematic of copper transport between the blood and brain in controls (A) and schizophrenia (B). Stars show copper. CTR1, white arrows; ATP7A, black arrows; ATP7B, striped arrows; BBB, blood brain barrier; BCB, blood cerebrospinal barrier. Thinner arrows indicate decreased protein levels.

Copper plays a key role in development and homeostatic function and is crucial for many cellular functions including monoamine metabolism, mitochondrial activity, and myelination (Sato et al. 1994). While copper and its enzymes are found outside of the brain, we will focus on brain. Copper dysfunction results in Wilson’s or Menkes disease, characterized by copper toxicity or deficiency, respectively (Wilson 1934; Menkes et al. 1962). Cellular copper is highly regulated, as free copper can induce oxidative stress and cellular damage (Halliwell and Gutteridge 2007). During normal function, copper is taken from the bloodstream across the blood brain barrier (BBB) into astrocytes and then neurons via CTR1 at the plasma membrane (Scheiber et al. 2010). Once inside the cell, copper is bound by metallochaperones (Maryon et al. 2013) and delivered to the trans-Golgi network (TGN). ATP7A is located within the TGN (Yamaguchi et al. 1996) and distributes copper to metallochaperones (e.g., SCO1) which transport the copper to their needed location within the cell (e.g., mitochondria)(Davies et al. 2013; Leary et al. 2007). CTR1 knockdown and/or loss results in developmental defects and lethality (Lee et al. 2001), and total loss of ATP7A results in Menkes disease and lethality (Menkes et al. 1962), exemplifying the incredible importance of these transporters in homeostasis and function.

Interestingly, decreasing copper activity through inhibited transporters or administering the copper chelator cuprizone to mice produces demyelination, altered neurotransmitters, and decreased oligodendrocytic protein expression (Gokhale et al. 2015; Gregg et al. 2009; Herring and Konradi 2011). Additionally, reduced copper activity results in schizophrenia-like behavioral impairments, such as deficits in novel object recognition, spatial memory tasks, pre-pulse inhibition, social interaction, and anxiety (Talbot et al. 2009; Gregg et al. 2009; Herring and Konradi 2011). The role of copper in schizophrenia has been rarely studied, but increased copper in the blood of schizophrenia subjects has been observed in many, but not all studies (see Vidovic et al. 2013 and references therein). However, the cause of this copper increase or resulting functional consequences have not been studied. We hypothesize that schizophrenia subjects exhibit excess blood copper due to faulty copper transport across the blood brain barrier (BBB).

Importantly, several intracellular components that interact with copper are reported to be abnormal in schizophrenia (Figure 1B). For example, schizophrenia subjects exhibit a deficit of SCO1, the metallochaperone responsible for delivering copper to the catalytic core of cytochrome C oxidase (COX) required for ATP synthesis (Leary et al. 2007; Purcell et al. 2014). Not only do schizophrenia subjects exhibit less COX protein, they also exhibit decreased ATP (Volz et al. 2000; Cavelier et al. 1995). Additionally, schizophrenia subjects exhibit decreased metallothionein and glutiothione, responsible for intracellular copper chaperoning and transport, that are rescued with antipsychotic treatment (Xuan et al. 2015; Do et al. 2000)(Figure 1B). While these deficits are well replicated, they have not been studied in the context of copper. Given that these mechanisms require copper, and that schizophrenia subjects exhibit alterations of intracellular copper pathway proteins (Figure 1B), copper may play an important role in schizophrenia pathology. Without proper regulation of the copper system and its transporters CTR1 and ATP7A, a cascade of schizophrenia-like effects can occur such as altered cellular energy metabolism (Leary et al. 2007; Volz et al. 2000; Cavelier et al. 1995), demyelination (Xuan et al. 2015), oxidative stress (Do et al. 2000), or even cellular death (Menkes et al. 1962; Halliwell and Gutteridge 2007; Lee et al. 2001).

The present study is the first to investigate dysbindin and copper transporters as a combined pathology in schizophrenia. The substantia nigra (SN) exhibits one of the highest levels of copper within the brain and the highest level of the copper chaperone Atox1 (Davies et al. 2013), indicating the SN has a high demand for cellular copper. Therefore, we measured 1) dysbindin isoforms 1A and 1B/C, encoded by risk factor gene *DTNBP1*; 2) copper transporter CTR1, responsible for copper transport across the BBB; 3) copper-transporting P-type ATPase, ATP7A, which works in conjunction with CTR1 for intracellular copper transport; and ATP7A homolog ATP7B, responsible for transfer of copper to the secretory pathway. We also conducted a preliminary analysis of medication status, as there are biological correlates to medication status. Our hypothesis is illustrated in Figure 1A. We hypothesize schizophrenia subjects exhibit deficits in copper transport in a copper-rich brain region potentially in relation to dysbindin alterations (Figure 1B). This work has been presented in preliminary form (Schoonover and Roberts 2016).

## Methods and Materials

### Postmortem Brains

Human brains were obtained from the Maryland Brain Collection with consent from the next of kin with IRB-approved protocols in accordance with all relevant guidelines and procedures. The work completed in this study was approved by the University of Alabama at Birmingham. The schizophrenia cohort was the same as previously studied (Schoonover et al. 2017). Schizophrenia cases (n=15) were matched and compared to normal controls (NC, n=11). As a preliminary investigation, the schizophrenia group was subdivided by treatment status: off medication (SZ-Off, n=4) or on medication (SZ-On, n=11). Diagnosis of schizophrenia was confirmed independently by two psychiatrists based on DSM criteria at the time of diagnosis (DSM-III-R through DSM-IV-TR) using the Structured Clinical Interview for the DSM (SCID). Subject clinical information (such as age of disease onset, symptomology, and treatment status) was obtained from autopsy and medical records, and family interviews. Placement into the off-medication schizophrenia group required being unmedicated for at least six months prior to death. Cases were selected based on the best match of demographic factors age, race, sex, postmortem interval (PMI), sample pH, and number of years frozen (Table 1A). Exclusionary criteria for this cohort were: history/evidence of intravenous drug abuse, HIV/AIDS, Hepatitis B, head trauma, comorbid neurological disorders, custodial death, fire victims, unknown next of kin, children, or decomposed subjects. In addition, comorbid mental illness in schizophrenia subjects and history of serious mental illness for NCs were exclusion factors. Agonal status has not been shown to have any effect on protein (Stan et al. 2006), and therefore is not an issue with the current investigation.

**Table 1.** (1A) Demographics and other information; mean and standard deviation. Some parameter information was not applicable for NCs. Therefore, an ANOVA was performed for parameters comparing three groups, and a t-test performed for parameters containing two groups. (1B) Correlational analyses of demographic and tissue quality markers. Comparisons between all proteins and demographic variables were made; significant p-values only are shown (bold). Only the relationship between dysbindin 1B/C and years frozen was significantly different between controls and schizophrenia subjects. While significant comparisons of coefficients were observed between medicated subjects for dysbindin 1B/C and postmortem interval, this demographic variable was not used as a covariate because no influence of medication status was found for dysbindin 1B/C. Abbreviations: Normal controls (NC); schizophrenia subjects (SZ); schizophrenia subject on medication (SZ-ON); schizophrenia subject off medication (SZ-OFF); postmortem interval (PMI); years sample has been frozen (YF); duration of illness (DUI); whole sample (WS); comparison of coefficients (CC); Caucasian (C); African American (AA).

| Table 1A. Demographics and Markers of Tissue Quality |  |  |  |  |  |  |  |  |  |
| --- | --- | --- | --- | --- | --- | --- | --- | --- | --- |
|  | # | Age, years | PMI, hours | pH | YF | Age of Onset | DUI, years | Race | Sex |
| NC | 11 | 50.9 ± 16.3 | 15.6 ± 7.1 | 6.7 ± 0.2 | 17.4 ± 4.7 | N/A | N/A | 8C/3AA | 9M/2F |
| SZ | 15 | 42.1 ± 12.2 | 13.6 ± 8.9 | 6.5 ± 0.4 | 19.3 ± 3.0 | 22.6 ± 5.1 | 18.6 ± 9.3 | 9C/6AA | 11M/4F |
| **t-test** | | p=0.13 | p=0.55 | p=0.30 | p=0.22 | N/A | N/A | $\chi^2=0.50$ | $\chi^2=0.6$ |
| SZ-On | 11 | 50.9 ± 16.3 | 15.2 ± 9.3 | 6.6 ± 0.3 | 18.1 ± 2.6 | 21.3 ± 4.8 | 19.9 ± 9.4 | 6C/5AA | 7M/4F |
| SZ-Off | 4 | 50.3 ± 17.5 | 11.0 ± 8.6 | 6.4 ± 0.4 | 21.8 ± 3.0 | 27.0 ± 4.2 | 14.0 ± 9.9 | 1C/3AA | 4M/0F |
| **ANOVA/**<br>**t-test** | | p=0.13 | p=0.45 | p=0.60 | p=0.15 | p=0.18 | p=0.46 | $\chi^2=0.20$ | $\chi^2=0.1$ |

| Table 1B. Demographic/Protein Correlations |  |  |  |  |  |
| --- | --- | --- | --- | --- | --- |
|  | N-ATP7A/<br>Age | ATP7B/<br>Age | T-CTR1/<br>YF | Dys 1BC/<br>YF | Dys 1BC/<br>PMI |
| **WS** | p=0.09<br>r= -0.34 | **p=0.04**<br>r= -0.41 | **p=0.03**<br>r= -0.43 | p=0.10<br>r= -0.34 | p=0.39<br>r= -0.18 |
| **NC** | **p=0.04**<br>r= -0.62 | p=0.19<br>r= -0.43 | p=0.17<br>r= -0.44 | **p=0.02**<br>r= -0.69 | p=0.73<br>r= 0.12 |
| **SZ** | p=0.51<br>r= -0.19 | p=0.77<br>r= -0.08 | p=0.35<br>r= -0.26 | p=0.62<br>r= 0.14 | p=0.39<br>r= 0.24 |
| **CC**<br>**NC vs SZ** | p=0.23 | p=0.41 | p=0.64 | **p=0.03** | p=0.79 |
| **OFF** | p=0.97<br>r= 0.03 | p=0.27<br>r= -0.73 | p=0.21<br>r= -0.79 | p=0.71<br>r= -0.29 | **p=0.03**<br>r= -0.97 |
| **ON** | p=0.92<br>r= -0.04 | p=0.16<br>r= -0.46 | p=0.82<br>r= 0.08 | p=0.38<br>r= 0.30 | p=0.45<br>r= 0.26 |
| **CC**<br>**OFF vs ON** | p=0.94 | p=0.68 | p=0.14 | p=0.57 | **p=0.03** |
| **CC**<br>**OFF vs NC** | p=0.47 | p=0.66 | p=0.57 | p=0.24 | **p=0.04** |
| **CC**<br>**ON vs NC** | p=0.16 | p=0.94 | p=0.25 | **p=0.02** | p=0.77 |

### Western Blotting

#### Tissue and Protein preparation

The SN was blocked as done previously (Schoonover et al. 2017). In SN coronal sections, caudal sections were caudal to third nerve rootlets. The blocks were trimmed to remove non-nigral tissue. A perimeter of approximately 2mm of non-nigral tissue remained. Frozen caudal SN was sonicated in lysis buffer (500ul/0.1 g of human tissue) containing Tris-HCL (pH 8.0), EDTA, sodium chloride, sodium dodecyl sulfate, and a protease inhibitor cocktail (Sigma; P8340). Tissue homogenate was centrifuged at 13,500 rpm for 15 mins at 4°C. Supernatant (total cell lysate) was then extracted and protein concentration determined via the Lowry method (Bio-Rad, Hercules, CA, USA; 500-0113, 500-0114).

#### Gel electrophoresis and western blotting

Western blots were used to measure N- and C-terminal ATP7A, ATP7B, extracellular and transmembrane CTR1, and dysbindin 1A and 1B/C protein levels and performed as previously reported (Schoonover et al. 2017), with the following exceptions. Samples intended for N-terminal ATP7A assays were not heated to avoid protein aggregation. Samples intended for all other assays were heated to 95°C for 5 mins. Protein extracts (60μg) were loaded onto 4%-20% gradient polyacrylamide gels (Lonza, Basel, Switzerland; 58505). Proteins were resolved by sodium dodecyl sulfate-polyacrylamide gel electrophoresis at 150V for 1 hr 15 mins, and then transferred at 30V for 21hrs onto polyvinylidene fluoride (PVDF) membranes (Bio-Rad, Hercules, CA, USA; 162-0174) at 4°C. Representative western blots are shown in Figure 2A; full blots are shown in Figure S1.

**Figure 2.**
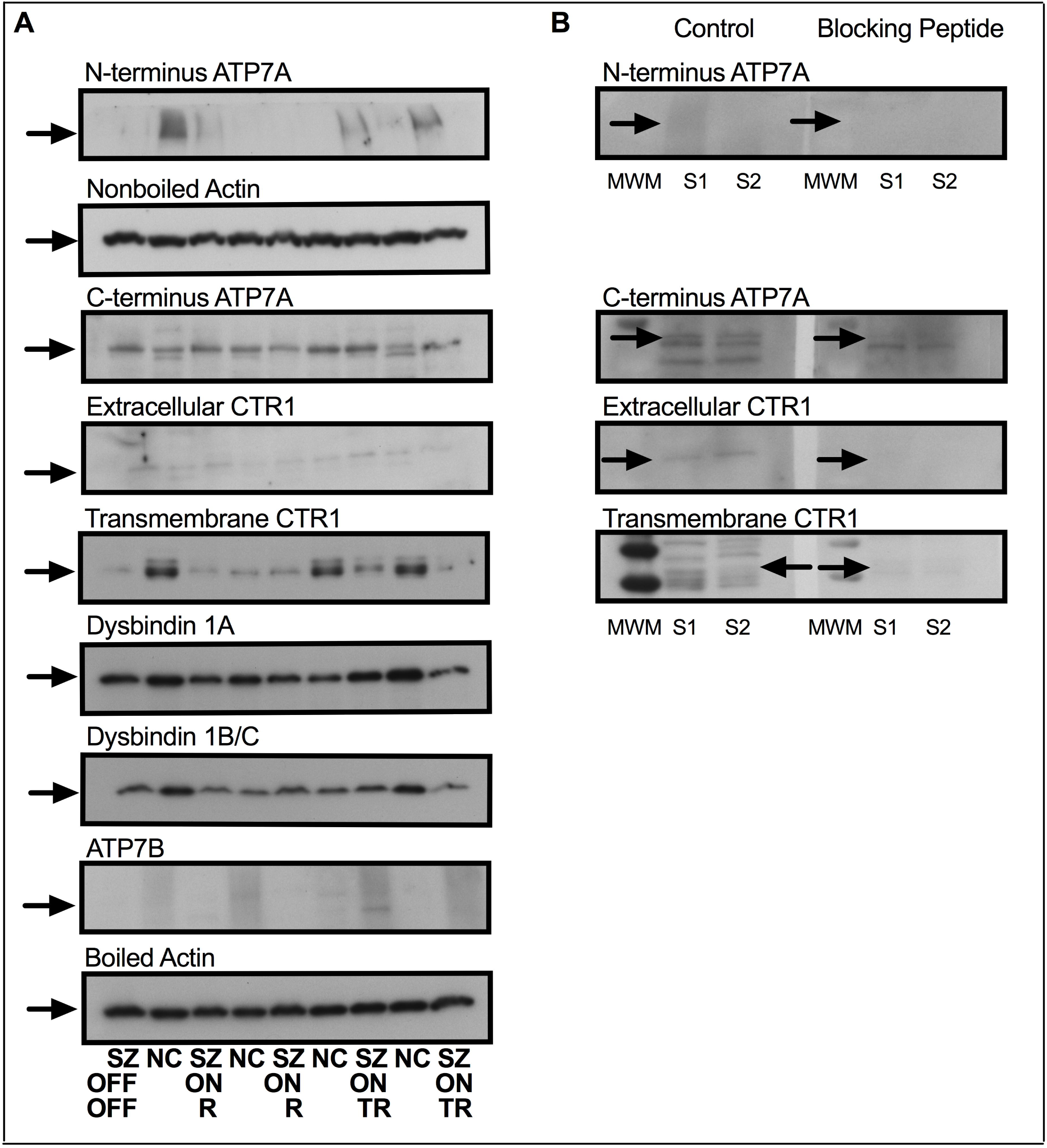
(A) Representative western blots. Arrows indicate the primary band at the expected molecular weight, with the exception of C-terminus ATP7A. Full blots with detailed molecular weight markers are shown in supplementary figure 1. Actin was a loading control. N-terminal ATP7A samples were not boiled; therefore non-boiled actin was the loading control. (B) Preadsorption blots for the proteins of interest. Arrows indicate the band that was measured and successfully blocked in two different normal controls (S1, S2). Control blots that incubated with primary antibody and no blocking peptide are shown on the left (control); preadsorbed blots are shown on the right (blocking peptide). MWM, molecular weight marker.

To target potential protein segment-dependent alterations of function, we used two antibodies with different epitopes for ATP7A and CTR1. The following antibodies and concentrations were used: rabbit anti-N-terminal ATP7A, 1:500, Novus Biologicals (NBP1-54906); rabbit anti-C-terminal ATP7A, 1:2000, Aviva Systems Biology (ARP33798_P050); rabbit anti-ATP7B, 1:1000, Abcam (ab135571); rabbit anti-transmembrane CTR1, 1:2,000, Novus Biologicals (NB100-402); rabbit anti-extracellular CTR1, 1:1000, Aviva Systems Biology (ARP43824_P050); rabbit anti-Dysbindin (targeting 1A and 1B/C), 1:2,000, Abcam (ab133652); and mouse anti-actin, 1:40,000, Millipore (MAB1501). Each blot contained a mixture of NC and schizophrenia subjects, and was performed in duplicate. The membranes were blocked for 1 hr in 5% milk in Tris-Buffered Saline with Tween 20 (TBST). Antigen presence was detected by incubating the primary antibody with the PVDF membrane 21hrs at 4°C. The bands were visualized using chemiluminescence (Bio-Rad; 170-5018), exposing Sigma-Aldrich Carestream Kodak BioMax XAR films (166-0760).

#### Experimental Controls

Some antibodies identified several bands; to determine the correct band to measure, we conducted a preadsorption experiment for N- and C-terminal ATP7A, and extracellular and transmembrane CTR1. Two identical blots containing samples from two separate controls were used. One blot was incubated with primary antibody; the second blot was incubated with a preadsorbed antibody mixture comprised of 1% milk containing primary antibody (the same concentration as used above) that incubated for 30 minutes at room temperature with the corresponding blocking peptide (1:500 concentration). The following blocking peptides were used: N-terminus ATP7A, Novus Biologicals (NBP1-54906PEP); C-terminus ATP7A, Aviva Systems Biology (AAP33798); transmembrane CTR1, Novus Biologicals (NB100-402PEP); and extracellular CTR1, Aviva Systems Biology (AAP43824). Verifying ATP7A and CTR1 antibody specificity with knockout animal models is not possible due to embryonic lethality (Menkes et al. 1962; Hua et al. 2010). Dysbindin knockout models are viable and well-characterized (Talbot 2009)-additionally, it was measured at its expected molecular weight in the present study.

### Analyses

#### Data

As described previously (Schoonover et al. 2017), films were scanned at 600 dpi using a flatbed scanner. Optical densities of the bands were measured using Image J-64 freeware (NIH). A step calibration tablet was used in order to create an optical density standard curve (Stouffer Industries Inc.; Mishawaka, IN, USA; T2120, series #130501) that each measurement was calibrated to. ImageJ was used to perform a background subtraction for each film. All optical density values for each protein were normalized to actin, and then to the averaged NC. These values were averaged for duplicate samples.

#### Statistics

Demographics and tissue quality were tested using ANOVA and/or t-tests. Categorical variables were assessed using a chi-square test. Outliers were detected using ROUT (Q=1.0%) and Grubb’s method (alpha=0.05) via Prism 7. Only one schizophrenia ATP7A value qualified for removal for the schizophrenia/normal control analysis. Two schizophrenia subjects had CTR1 values that qualified for removal for the schizophrenia/normal control analysis and the treatment status analysis. The data were assessed for normality with both a Shapiro-Wilk test and a D’Agostini and Pearson test. If both tests revealed normally distributed data, parametric tests were used (two groups, unpaired t-test; three groups, one-way ANOVA). If not, non-parametric tests were used (two groups, Mann-Whitney U test; three groups, Kruskal-Wallis H test).

Significant omnibus tests were followed by planned, uncorrected comparisons suggested by Prism 7: parametric tests were followed by an uncorrected Fisher’s LSD test, while nonparametric tests were followed by an uncorrected Dunn’s comparison. Furthermore, Brown-Forsythe and Bartlett’s tests were used to compare standard deviations. Correlational analyses were performed between results and PMI, age, years frozen, and pH to elucidate potential relationships among these variables. Based on these results, years frozen was used as a covariate in the dysbindin 1B/C analysis.

## Results

### Demographics

The control (NC) and schizophrenia groups (as a whole and when subdivided by treatment status) were well matched for demographic and tissue quality variables and did not significantly differ (Table 1A). Correlation analysis between protein concentrations and demographic variables only showed a significant between-group difference for dysbindin 1B/C and years frozen (Table 1B). Thus, years frozen was used as a covariate in the dysbindin 1B/C analysis.

### Protein Alterations

All proteins, with the exception of C-terminus ATP7A and transmembrane CTR1, were measured at their expected molecular weights and were successfully blocked by their respective blocking peptide (Figure 2A,B; for full blots see Supplementary Figure S1). While not at their expected molecular weight, the measured C-terminus ATP7A and transmembrane CTR1 bands were verified with blocking peptide.

While levels of the ATP7A N-terminus protein were not different between groups, schizophrenia subjects exhibited significantly more C-terminus ATP7A protein than controls (p=0.005; Figure 3A). However, when subdivided by medication status, the N-terminus protein levels were significantly lower in unmedicated subjects versus medicated subjects (p=0.007) and controls (p=0.017; Figure 3B). Analysis of treatment status revealed that C-terminal ATP7A protein levels were significantly increased only in medicated subjects (p=0.013; Figure 3B).

**Figure 3.**
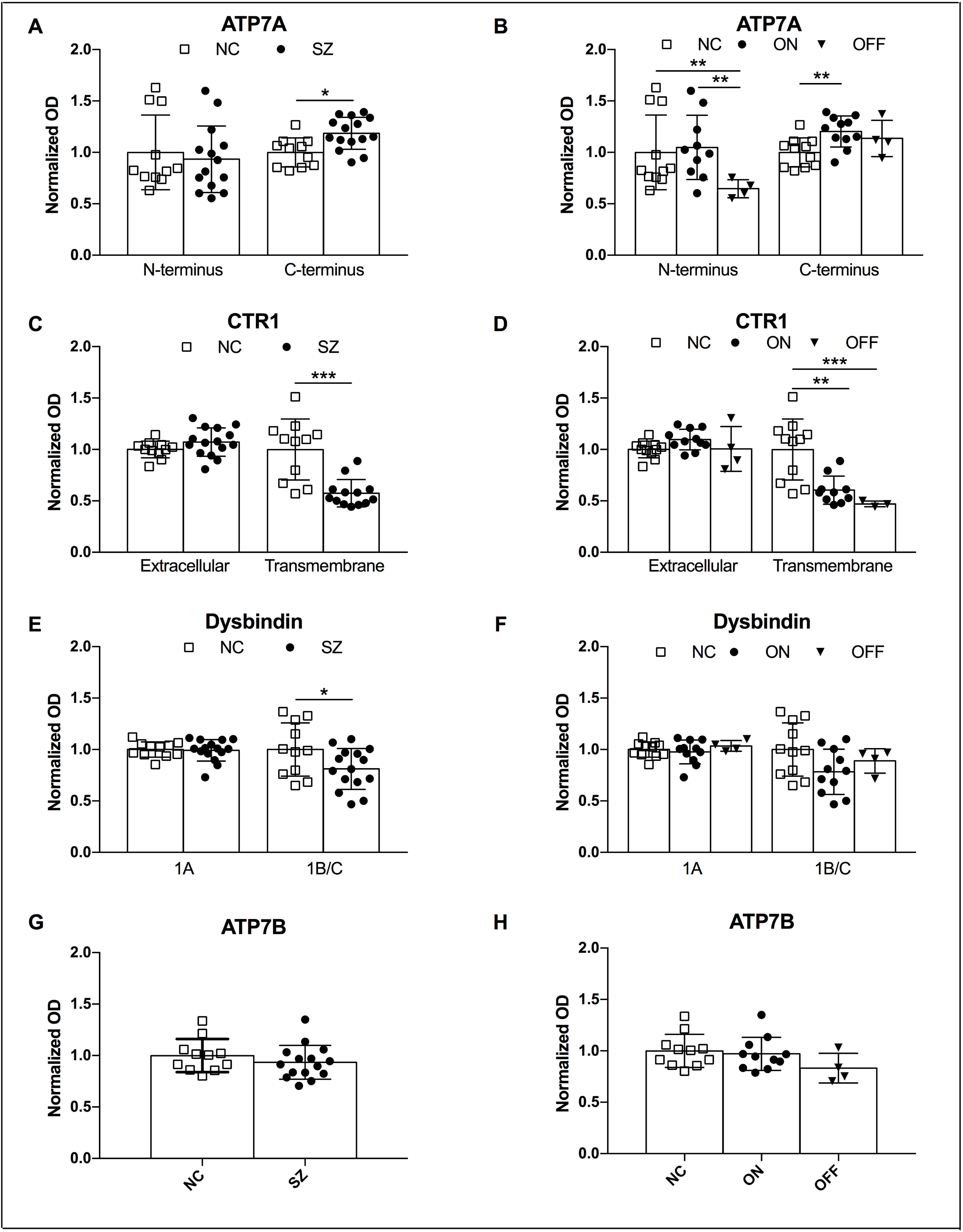
Protein levels of ATP7A (A-B), CTR1 (C-D), dysbindin 1A, 1B/C (E-F) and ATP7B (G-H) for analysis of diagnostic group and treatment status. Error bars represent standard deviation. Significant omnibus ANOVA results are: N-terminal ATP7A: p=0.02; C-terminal ATP7A: p=0.01; transmembrane CTR1: p=0.001. Significant ANOVAs were followed by post hoc tests illustrated by the following: *: p<0.05; **: p<0.01; ***: p<0.001.

Schizophrenia subjects had significantly lower levels of transmembrane CTR1 than did controls (p=0.0003), but exhibited no changes in the extracellular component of the protein (Figure 3C). Analysis of treatment status revealed no antipsychotic effects. The transmembrane CTR1 was decreased by a similar amount in both medicated (p=0.007) and unmedicated schizophrenia patients (p=0.001; Figure 3D), and there was no effect of medication on the extracellular CTR1 levels.

Dysbindin 1A protein levels were not altered in the whole schizophrenia group or when divided by treatment status (Figure 3E, 3F). Decreased dysbindin 1B/C was observed in schizophrenia subjects versus controls (p=0.046; Figure 3E), but no effects of antipsychotics were observed (Figure 3F). However, the significant effect was reduced to a trend when the variable years frozen was implemented as a covariate (p=0.062).

No changes of ATP7B were observed in the whole schizophrenia group or when subdivided by treatment status (Figure 3G; Figure 3H).

Schizophrenia subjects treated with antipsychotic medication were further divided by antipsychotic type (typical versus atypical antipsychotics). No differences were found (data not shown).

## Discussion

This is the first study to investigate dysbindin and copper transporters as a combined pathology in schizophrenia. Taken together, we observed protein region-specific alterations of ATP7A and CTR1, some of which appear to be modulated by antipsychotic treatment. Additionally, we observed isoform-specific decreases of dysbindin. These results suggest that copper homeostasis is altered in schizophrenia, and merit further study.

### Limitations

Our sample size was small and as is typical for postmortem investigations, none of our subjects were first-episode and/or antipsychotic-naïve (Schoonover et al. 2017; Howes et al. 2013). Medication history could affect our results, so we divided the schizophrenia group by treatment status and type to eliminate this confounding variable. However, there were only four unmedicated cases, and therefore analysis of treatment status can only be considered preliminary. No differences were found between schizophrenia subjects treated with typical versus atypical antipsychotics (data not shown).

### Isoform-Specific Dysbindin Alterations

Given the association between dysbindin allelic variations and protein expression in schizophrenia (Talbot 2009; Straub et al. 2002; Tang et al. 2009; Weickert et al. 2004, 2008), we anticipated decreased dysbindin expression in schizophrenia. Interestingly, schizophrenia subjects exhibited decreased dysbindin 1B/C, but not isoform 1A. Dysbindin 1 has three isoforms known as dysbindin 1A, 1B, and 1C (Talbot et al. 2009; Talbot et al. 2011). Dysbindin 1 A is found in the post-synaptic density and thought to be involved in dendritic spine homeostasis, while dysbindin 1B is found exclusively in synaptic vesicles in the presynaptic axon terminal (Talbot et al. 2011), and is involved in glutamatergic transmission (Numakawa et al. 2004). Dysbindin 1C is found in both places, but primarily is located in the post-synaptic density and involved in dendritic spine function (Talbot et al. 2011). The substantia nigra is relatively spine free (Schwyn and Fox 1974) and the majority of synapses are GABAergic (Tepper and Lee 2007). Approximately 30% of inputs to the SN are glutamatergic (Smith et al. 1996, Parent et al. 1999). Our findings of modest decreases in the dysbindin 1B/C isoform implicate abnormalities in presynaptic glutamate terminals, which can lead to impaired glutamate transmission (Numakawa et al. 2004). Since neurons in the substantia nigra are mostly aspiny, the isoforms 1A and 1C, which are involved in spine function, may have little consequence in this brain region.

Other groups have also found decreases in specific isoforms in schizophrenia in various brain regions, such as superior temporal gyrus (1A), dorsolateral prefrontal cortex (1C), and hippocampus (1B/C) (Tang et al. 2009; Talbot et al. 2011). These results, taken together, indicate isoform-specific roles of dysbindin in schizophrenia that are brain region-specific. Given that dysbindin isoforms are differentially associated with synaptic function (Talbot et al. 2009; Numakawa et al. 2004), which is repeatedly found to be abnormal in schizophrenia (Faludi and Mirnics 2011; Roberts et al. 2012), isoform-specific changes observed in the present study are consistent with previous findings.

### Copper Transporter Deficits

Based on a report of decreased copper transporters in dysbindin knockout mice (Gokhale et al. 2015), we hypothesized dysregulated copper transporters CTR1 and ATP7A as a consequence of downregulated dysbindin in schizophrenia subjects. While our results support our hypothesis, we did not anticipate the protein-region specific alterations we observed. We observed a surplus of C-, but not N-terminus, ATP7A in the whole schizophrenia group. The surplus of C terminal ATP7A was significant only in medicated subjects; however, there was no difference between medicated and unmedicated subjects, suggesting no effect of antipsychotic drugs. However, analysis of treatment status revealed what may be an inherent deficit of N-terminal ATP7A in schizophrenia that is rectified with antipsychotic treatment.

While protein region differences ATP7A may initially seem puzzling (given that ATP7A was measured at its expected molecular weight), it potentially highlights a recurring and well-established alteration of protein phosphorylation in schizophrenia. For example, dysregulated kinase activity has been observed (Banerjee et al. 2015; Singh 2013), affecting, for example, phosphorylation of MAP2 and NMDAR proteins (Shelton et al. 2015; Banerjee et al. 2015; Ramkumar et al. 2018). The antibody used to assess N-terminal ATP7A does not recognize phosphorylated ATP7A protein. Therefore, perhaps the decreased N-terminal ATP7A observed in unmedicated schizophrenia subjects does not represent a loss of protein, but rather post-translational modifications that alter the phosphorylation state. In fact, upon the binding of copper to one of the six metal binding domains (Figure 4A) of N-terminal ATP7A, γ-phosphate is transferred from ATP to the Asp residue within the P domain of the protein (see Figure 4 in Lutsenko et al. 2007; Valerde et al. 2008). This results in transient phosphorylation that is reversed (dephosphorylated) upon the release of copper into the lumen (Voskoboinik et al. 2001; Valverde et al. 2008).

**Figure 4.**
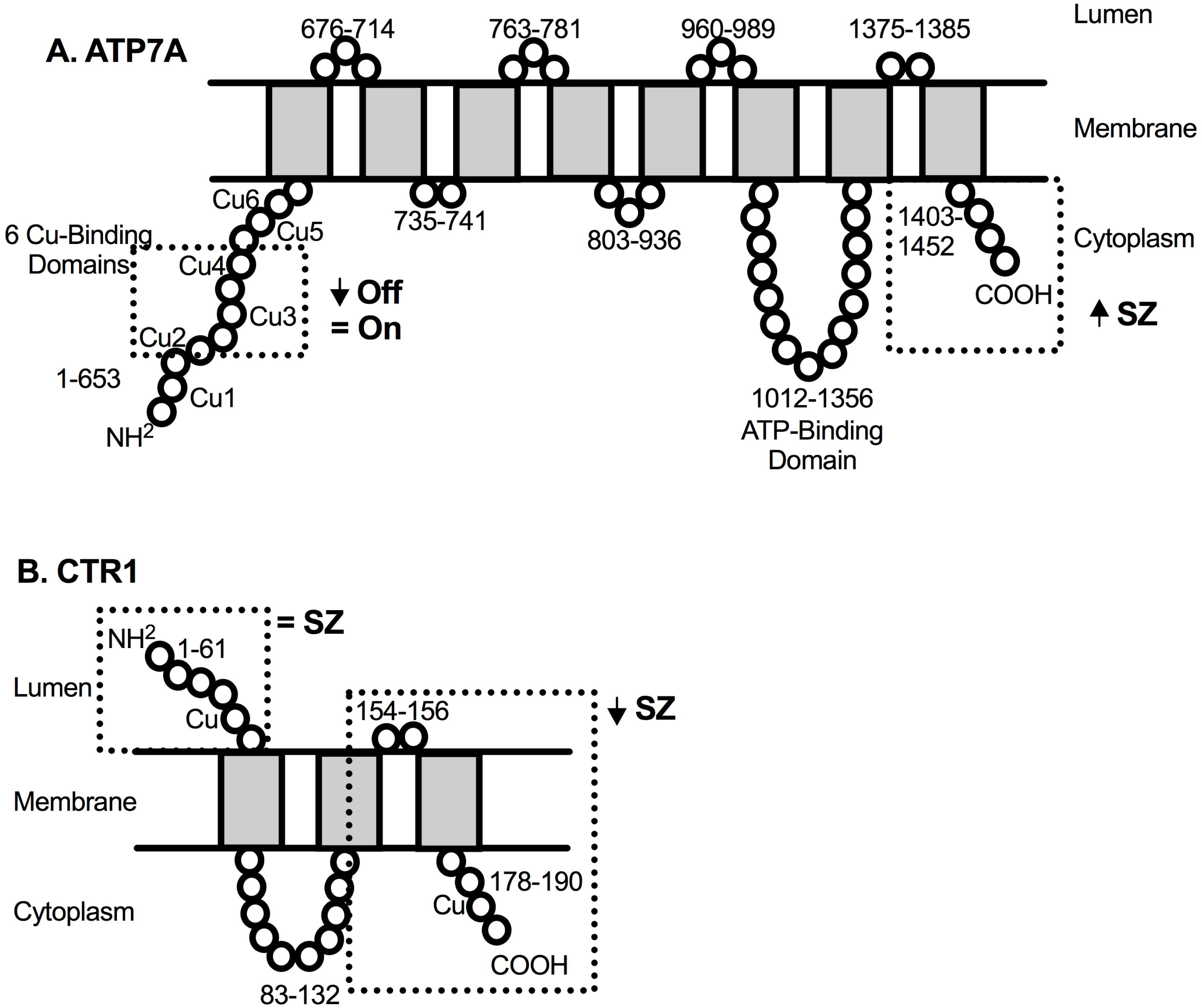
Numerical labeling of amino acid ranges are shown for each protein segment. Dotted boxes indicate antibody-specific epitopes. **A.** ATP7A protein structure. N-terminal antibody specifically binds to 225-273aa; C-terminal antibody binds to 1403-1452aa. **B.** CTR1 protein structure. Extracellular CTR1 antibody binds to 19-68aa; transmembrane CTR1 antibody specifically binds to 140-190aa. **C.** Intracellular diagram of copper chaperones and enzymes and schizophrenia-related alterations. Copper enters neurons after transport from astrocytes via CTR1 and is immediately bound by MT/<u>GSH. MT/GSH</u> then delivers copper to chaperone ATOX1 or the TGN where it is delivered to other metalloenzymes (e.g., SCO1 → COX) via ATP7A. The SN exhibits more ATOX1 than other brain regions. SCO1 is a copper-requiring enzyme involved in the last step of the electron transport chain of ATP synthesis. SCO1, MT/GSH, COX, ATP, ATP7A and CTR1 are all downregulated or altered in schizophrenia. Abbreviations: MT, metallothionein; GSH, glutathione; APD, antipsychotic drug; SN, substantia nigra; COX, cytochrome c oxidase. References: 1) The present paper; 2) Scheiber et al. 2010; 3) Maryon et al. 2013; 4) Do et al. 2000; 5) Xuan et al. 2015; 6) Davies et al. 2013; Prohaska 1987; 7) Petris et al. 1996; 8) Leary et al. 2007; 9) Purcell et al. 2014; 10) Cavelier et al. 1995; and 11) Volz et al. 2000.

Like ATP7A, we also observed domain-specific alterations of CTR1 in schizophrenia. Specifically, the transmembrane and intracellular segment of CTR1 was decreased, but not the extracellular portion (Figure 4B). CTR1, like ATP7A, is a transmembrane protein. The extracellular and intracellular portions both possess copper-binding domains (Eisses and Kaplan 2005); knockout of either region impairs copper uptake, though <40% of uptake remains functional (Eisses and Kaplan 2005). Our transmembrane protein analysis includes the extracellular pore/channel opening (Eisses and Kaplan 2005); deficits in this region may affect extracellular copper binding and its transport to the cytosol, potentially resulting in decreased copper transport into the cell. The mechanism behind domain-specific alterations of CTR1 in schizophrenia remains untested. However, perhaps CTR1 transcription regulation is pathologically altered. In yeast, *CTR1* contains a copper response element (CuRE) in its promoter, targeted by transcription factor Mac1, which regulates CTR1 protein expression in a copper-dependent manner (Labbe et al. 1997). During times of copper starvation, *CTR1* should be upregulated. However, single or multiple point mutations within the CuREs suppress both copper-dependent repression and expression of *CTR1* (Labbe et al. 1997). Given the plethora of genetic alterations in schizophrenia, perhaps CTR1 is yet another to add to the list. Additionally, CTR1 may contribute to the genetic alterations in schizophrenia, as it is required for Ras and MAPK signaling (Turski et al. 2012; Tsai et al. 2012). Further study is needed to determine genetic regulation of *CTR1* expression in human cells and its implications for schizophrenia.

Given the decreased CTR1, buildup of copper in the plasma of schizophrenia subjects (Vidovic et al. 2013), and the dysregulation of kinases and phosphorylation in schizophrenia (Banerjee et al. 2015; Sing 2013; Shelton et al. 2015; Ramkumar et al. 2018), we suggest a potential alteration of the copper-sensing system of the metal binding domains and the cascading system of phosphorylation thereafter in unmedicated schizophrenia that is potentially corrected with antipsychotic treatment. Our finding of increased C-terminal ATP7A in medicated subjects suggests further efforts by the cell to restore copper homeostasis. ATP7A is constantly recycled between the plasma membrane and the TGN, but maintains a steady-state localization in the TGN during normal copper conditions (Yamaguchi et al. 1993). The carboxy terminus of the protein is responsible for endocytosis from the plasma membrane to the TGN (Petris et al. 1996; Petris and Mercer 1999; Francis et al. 1999). By increasing localization of ATP7A to the TGN, relatively increased copper insertion into metalloenzymes and the secretory pathway should occur.

In brain, ATP7B moves copper from the parenchyma through the blood brain barrier to the blood (Figure 1A). ATP7B deficits result in Wilson’s disease, characterized by high copper accumulation in brain and liver and low accumulation in blood (Wilson 1934). It is therefore possible that an increase in ATP7B could be causing the increases in blood copper levels often reported in schizophrenia. We did not observe any alterations of the copper transporter ATP7B, narrowing down the possible mechanisms by which increased blood copper in schizophrenia could occur.

### Implications

Given that ATP7A, dysbindin and CTR1 are located, in part, in the astrocytic end feet that form the BBB (Scheiber et al. 2010; Iijima et al. 2009; Nyguyen et al. 2017) our results suggest abnormalities of the BBB in schizophrenia subjects. If less copper is transported across the BBB in schizophrenia, excess blood copper and a brain copper deficit could result. These results could explain the conundrum of elevated copper levels in the blood of patients (see Vidovic et al. 2013 and references therein), and the pathologies and behaviors reminiscent of schizophrenia that occur during cellular copper starvation (Gokhale et al. 2015; Gregg et al. 2009; Herring and Konradi 2011). In mice, copper starvation results in demyelination, impaired prefrontal cortex function, and schizophrenia-like behaviors such as deficits in novel object recognition, spatial memory tasks, pre-pulse inhibition, social interaction, and anxiety (Talbot et al. 2009; Gregg et al. 2009; Herring and Konradi 2011). Therefore, brain copper starvation may be an underlying contributor to the cognitive symptoms of schizophrenia.

### Conclusions

The current study provides the first evidence of abnormal brain copper homeostasis in schizophrenia. Our findings suggest protein-region specific abnormalities in copper transport and isoform-specific dysbindin alterations. Intracellular copper binding appears to be decreased while ATP7A endocytosis mediation is increased, but only the binding is modulated by antipsychotic treatment. Our results suggest extracellular copper binding and transport into the cell via CTR1 is impaired in schizophrenia, and not rescued with treatment. Our findings not only provide information on a potentially new pathological mechanism, but may provide a link between several well-known but seemingly segregated findings in schizophrenia. Lastly, these results may elucidate the paradox of excess copper in schizophrenia blood and the schizophrenia-like pathology that occurs in a copper-deficit state. However, schizophrenia is a complex disorder and therefore copper alterations as a contributing pathology in schizophrenia merit further study. Further elucidation could provide vital information about the cellular pathology of schizophrenia and open new avenues for treatment.

## Acknowledgements

We would like to thank the Maryland Brain Collection staff for the samples used in this study. Additionally, this work was supported by the National Institute of Mental Health R0166123 and R21MH108867 to R.C.R, as well as the National Institute of Neurological Disorders and Stroke F99NS105208 to K.E.S.

